# A Patient-Specific Electrical Twin of Intracranial Pressure Dynamics Validated by Clinical Infusion Tests

**DOI:** 10.64898/2026.05.17.725750

**Authors:** Leszek Herbowski

## Abstract

Understanding intracranial pressure (ICP) dynamics is essential for interpreting clinical infusion tests used in the diagnosis of cerebrospinal fluid circulation disorders. However, the complex coupling between vascular pulsations, cerebrospinal fluid flow, and intracranial compliance makes quantitative interpretation of these tests challenging.

Here, I present a patient specific simulation framework based on an extended electrical analog model that reproduces intracranial pressure dynamics observed during clinical infusion tests. The model integrates physiological inputs including arterial blood pressure, heart rate, respiratory rhythm, and resistance to cerebrospinal fluid outflow derived from clinical data. Built upon the classical Ursino framework, the model incorporates several modifications enabling realistic representation of physiological pulsations and infusion test conditions. The resulting system functions as a hybrid electrical–numerical simulation model representing a simplified digital electrical twin of intracranial hydrodynamics.

The model was validated using data from 21 clinical infusion tests performed in patients with suspected normal pressure hydrocephalus. Simulated intracranial pressure recordings were compared with clinical measurements using regression and residual analysis. The simulations demonstrated strong agreement with measured data, with a mean correlation coefficient of r = 0.95 (95% CI 0.94 - 0.96), mean residual values within -1.71 to +1.68 mmHg, and a mean root mean square error (RMSE) of 2.07 mmHg.

These results demonstrate that the proposed model accurately reproduces the dynamic behavior of intracranial pressure observed during clinical infusion tests. The framework provides a physiologically grounded computational tool for studying patient specific intracranial dynamics and may support improved interpretation of infusion test results in clinical practice.

## 1. Introduction

The concept of using analogies to understand physical phenomena has a long tradition in science. In the late nineteenth century, electrical phenomena were often explained by comparison with fluid flow in pipes, an approach known as the “drain-pipe theory.” Interestingly, several decades later this analogy was reversed: electrical circuits began to be used to describe biological flow systems such as blood circulation and cerebrospinal fluid dynamics.

The concept of cerebrospinal fluid circulation, sometimes referred to as the “third circulation,” was introduced by Cushing in the early twentieth century [1]. Understanding the hydrodynamic behavior of this system has since become crucial for interpreting pathological conditions such as hydrocephalus and other disorders affecting intracranial pressure dynamics.

Early attempts to model intracranial dynamics relied on simplified compartmental representations in which the intracranial system was treated as one or several interacting compartments characterized by physical parameters such as compliance and resistance. The pioneering work of Marmarou introduced nonlinear mathematical models describing cerebrospinal fluid dynamics [2,3]. Subsequently, Ursino developed a comprehensive multi-compartment electrical analog model describing interactions between cerebral blood flow, intracranial pressure, and cerebrospinal fluid circulation [4–9]. Later simplifications of this model further improved its practical applicability [10].

Another early contribution to the electrical modeling of cerebrospinal fluid dynamics was provided by Agarwal and colleagues, who proposed a lumped-parameter model of intracranial compartments in the 1960s [12]. Their work can be considered one of the earliest electrical analog representations of the intracranial system.

Although modern neuroimaging techniques allow increasingly detailed visualization of cerebrospinal fluid dynamics, they have not yet established a universal gold standard for diagnosing disorders such as normal pressure hydrocephalus (NPH) [13–15]. In clinical practice, the infusion test remains an important diagnostic method for evaluating cerebrospinal fluid dynamics and resistance to CSF outflow [16–18].

Electrical analog models of intracranial dynamics provide an intuitive framework for integrating multiple physiological processes. However, many previously described models remain primarily theoretical and rarely incorporate patient-specific physiological inputs or direct validation using infusion tests.

The aim of this study was twofold. First, to analyze limitations of existing electrical analog models of intracranial dynamics and introduce necessary modifications improving their physiological realism. Second, to implement a patient-specific simulation framework capable of reproducing intracranial pressure changes observed during infusion tests. The overall methodological framework of the study, including model reconstruction and patient-specific simulation workflow, is illustrated in Fig. 1.

**Fig. 1.**
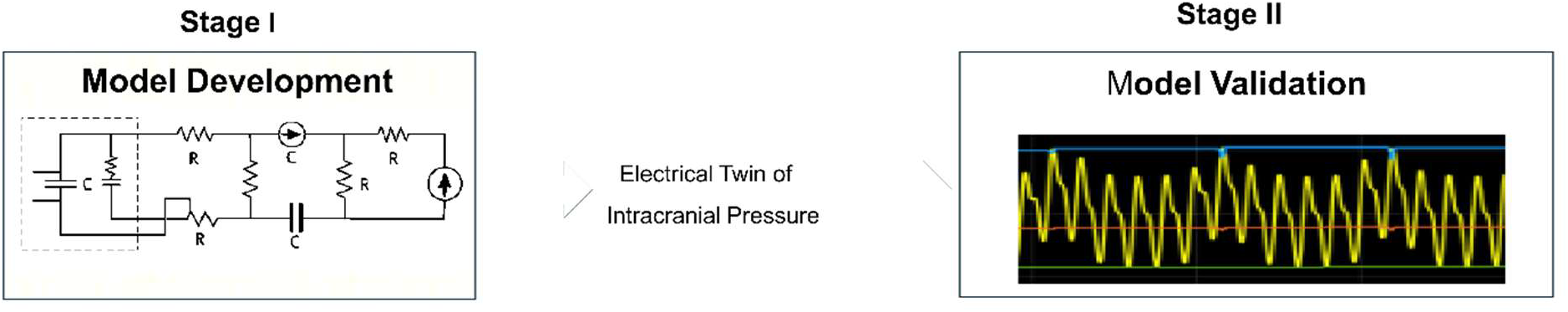
Study workflow illustrating the development and validation of the electrical analog model of intracranial pressure dynamics. The diagram presents the two-stage methodology used in this study. Stage I involves reconstruction and modification of the electrical analog model of intracranial dynamics based on previously published frameworks. Stage II includes patient-specific simulations of intracranial pressure dynamics and validation of the model using clinical infusion test recordings.

By integrating electrical analog modeling with numerical simulation in MATLAB/Simulink and clinical data from infusion tests, the resulting framework can be considered a hybrid electrical-numerical simulation model representing a simplified digital “electrical twin” of intracranial pressure dynamics.

## 2. Materials and Methods

### 2.1 General data

The methodological framework illustrated in Fig. 1 was implemented using MATLAB (version R2023a) and Simulink software (MathWorks Inc., Natick, MA, USA). Intracranial pressure simulations were performed using an electrical analog model composed of resistors and capacitors representing physiological components of the intracranial system.

The model was constructed in Simulink using block-based electrical elements. The software automatically transformed the circuit structure into a system of differential equations solved numerically during simulation.

Because several model parameters are dynamic (for example, variable compliance and autoregulatory mechanisms), an initial stabilization phase was required to allow the system to reach steady-state conditions before the infusion test simulation. Therefore, a 30-minute baseline period was introduced prior to the simulated infusion phase.

The study design consisted of reconstructing the electrical analog model based on the Ursino framework with modifications described in previous studies. If the reconstructed model failed to reproduce physiologically plausible ICP dynamics, further modifications were introduced. Patient-specific physiological inputs included systolic and diastolic arterial blood pressure, heart rate, respiratory rate, and resistance to cerebrospinal fluid outflow (R_o_). These parameters were derived from clinical infusion tests performed in patients suspected of normal pressure hydrocephalus.

### 2.2 Ethics statement

All infusion tests were performed as part of routine clinical management. The study was approved by the regional ethics committee and conducted in accordance with the Declaration of Helsinki.

### 2.3 Statistical analysis

Statistical analysis was performed to evaluate the agreement between simulated and clinically measured intracranial pressure (ICP) values obtained during infusion tests. Simulated ICP time series were temporally aligned with the corresponding clinical recordings, and point-to-point comparisons were performed across the entire duration of each infusion test. Each simulated ICP value was paired with the corresponding measured ICP value at the same time point. The agreement between simulated and measured ICP values was assessed using linear regression analysis and Pearson correlation coefficient (r). The coefficient of determination (R^2^) was calculated to quantify the proportion of variance in the measured ICP explained by the model. Residual analysis was performed by calculating the difference between simulated and measured ICP values at each time point. For each infusion test, the minimum and maximum residual values were determined to define the residual range. The mean residual range across all cases was also calculated. The root-mean-square error (RMSE) was computed as a measure of the overall deviation between simulated and measured ICP values.

Confidence intervals (95% CI) for the correlation coefficients were estimated to assess the robustness of the observed relationships. Statistical significance was defined as p < 0.05. All statistical analyses were performed using MATLAB (MathWorks Inc., Natick, MA, USA) and Statistica software (version 10, StatSoft Inc., Tulsa, OK, USA).

### 2.4 Extended electrical analog and input signals

An additional resistance element (R_t_) was introduced between the arterial and cerebrospinal fluid compartments to attenuate arterial pulsations transmitted to the CSF space and to prevent non-physiological amplification of intracranial pressure pulsatility during infusion test simulations. The conceptual structure of the extended electrical analog model is shown in Fig. 2, while the full implementation of the model in MATLAB/Simulink is provided in Supplementary Fig. S1.

**Fig. 2.**
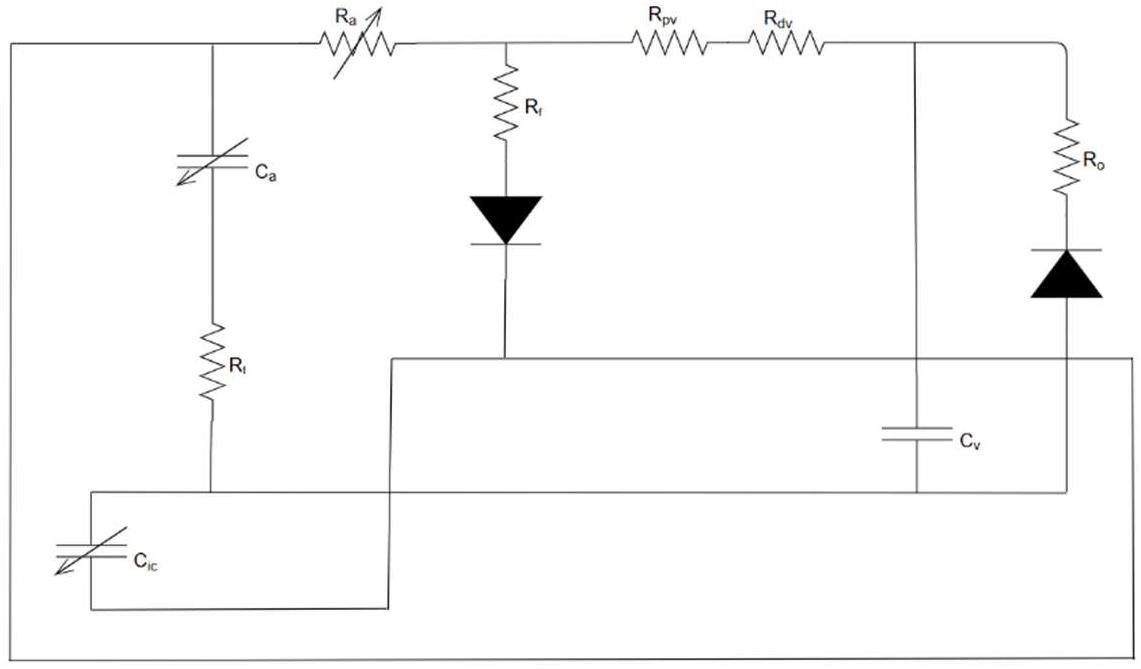
Electrical analog model of intracranial dynamics. Simplified conceptual diagram of the extended lumped-parameter electrical analog representing intracranial pressure dynamics. The model integrates arterial, venous, cerebrospinal fluid (CSF), and spinal compartments using electrical elements such as resistors, capacitors, and controlled sources. The diagram illustrates the principal pathways governing pressure–flow interactions within the intracranial system. The full model implementation is provided in Supplementary Fig. S1.

The electrical model was further extended with modules enabling the introduction of patient-specific physiological signals and perturbations corresponding to infusion tests.

Three main physiological input modules were implemented in the model:

#### Heart rate module

This module generates pulsatile signals corresponding to cardiac activity, allowing the simulation of arterial pressure oscillations.

#### Pulsatile arterial pressure module

This module introduces systolic and diastolic pressure variations reflecting physiological arterial blood pressure waveforms.

#### Respiratory-venous module

Respiratory oscillations influence venous pressure within the intracranial venous sinuses. Based on physiological observations, inspiratory phases were modeled as shorter and more pronounced than expiratory phases. The resulting venous sinus pulsations were implemented as periodic pressure fluctuations.

The mean venous sinus pressure was calculated using the relationship:

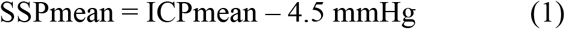

as suggested by previous physiological studies [19-21].

### 2.5 Infusion test simulation module

A dedicated simulation module reproduced the clinical infusion test. The module introduced fluid into the cerebrospinal fluid compartment at a constant rate corresponding to the clinical procedure.

The duration of simulated infusion tests matched the duration of the clinical recordings, including baseline, infusion, plateau, and recovery phases.

## 3. Results

### 3.1 Patient characteristics

Twenty-one patients (13 women and 8 men) with a mean age of 55 years were included in the study. A total of 21 infusion tests were analyzed (20 intraventricular and 1 lumbar).

Mean baseline intracranial pressure was 11.4 mmHg and mean resistance to CSF outflow was 12 mmHg/(ml/min). Detailed demographic and physiological parameters are presented in Table 1.

**Table 1.**
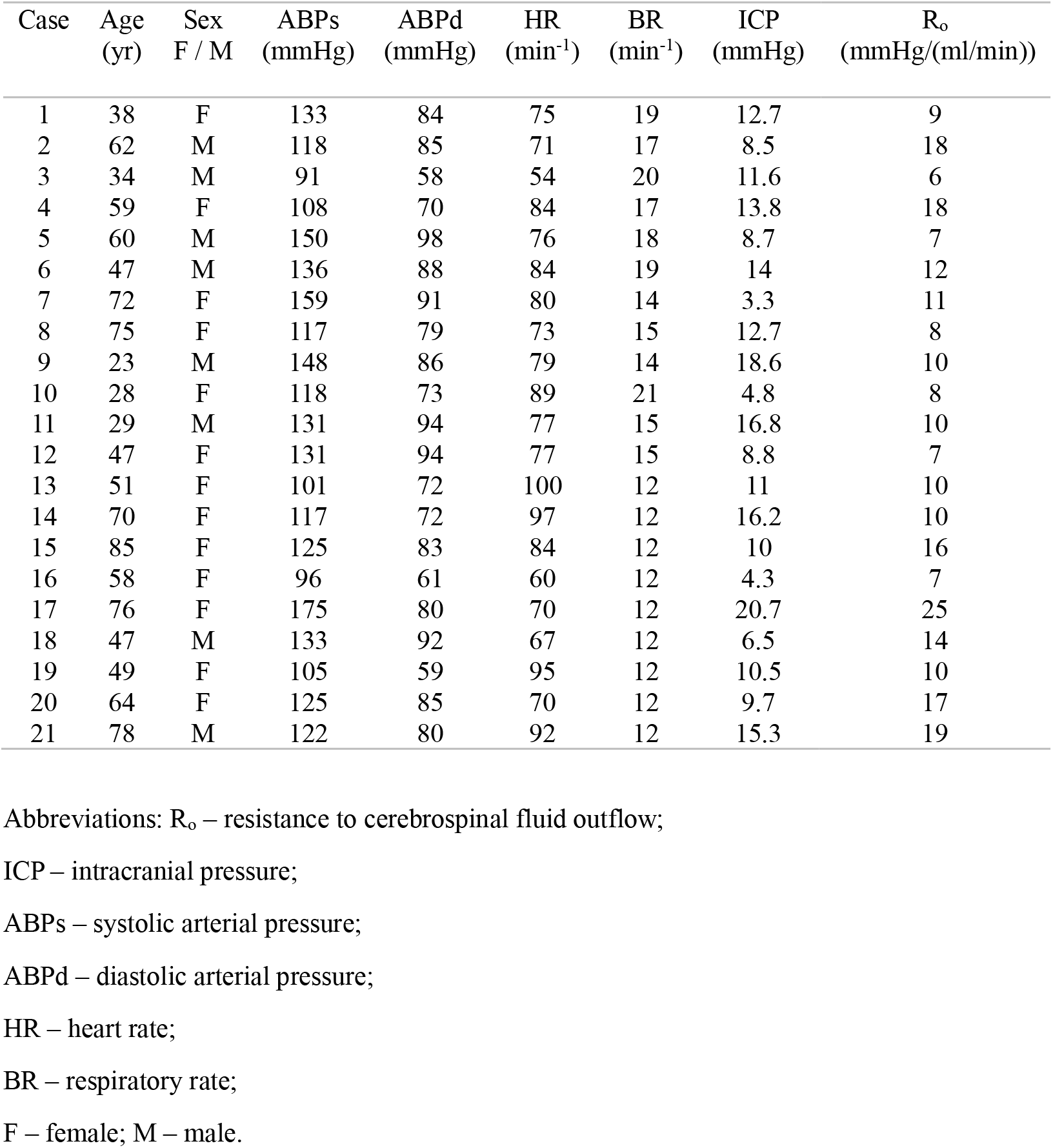
Patient-specific physiological parameters used as input signals for the infusion test simulations.

| Case | Age<br>(yr) | Sex<br>F / M | ABPs<br>(mmHg) | ABPd<br>(mmHg) | HR<br>(min<sup>-1</sup>) | BR<br>(min<sup>-1</sup>) | ICP<br>(mmHg) | R<sub>o</sub><br>(mmHg/(ml/min)) |
| --- | --- | --- | --- | --- | --- | --- | --- | --- |
| 1 | 38 | F | 133 | 84 | 75 | 19 | 12.7 | 9 |
| 2 | 62 | M | 118 | 85 | 71 | 17 | 8.5 | 18 |
| 3 | 34 | M | 91 | 58 | 54 | 20 | 11.6 | 6 |
| 4 | 59 | F | 108 | 70 | 84 | 17 | 13.8 | 18 |
| 5 | 60 | M | 150 | 98 | 76 | 18 | 8.7 | 7 |
| 6 | 47 | M | 136 | 88 | 84 | 19 | 14 | 12 |
| 7 | 72 | F | 159 | 91 | 80 | 14 | 3.3 | 11 |
| 8 | 75 | F | 117 | 79 | 73 | 15 | 12.7 | 8 |
| 9 | 23 | M | 148 | 86 | 79 | 14 | 18.6 | 10 |
| 10 | 28 | F | 118 | 73 | 89 | 21 | 4.8 | 8 |
| 11 | 29 | M | 131 | 94 | 77 | 15 | 16.8 | 10 |
| 12 | 47 | F | 131 | 94 | 77 | 15 | 8.8 | 7 |
| 13 | 51 | F | 101 | 72 | 100 | 12 | 11 | 10 |
| 14 | 70 | F | 117 | 72 | 97 | 12 | 16.2 | 10 |
| 15 | 85 | F | 125 | 83 | 84 | 12 | 10 | 16 |
| 16 | 58 | F | 96 | 61 | 60 | 12 | 4.3 | 7 |
| 17 | 76 | F | 175 | 80 | 70 | 12 | 20.7 | 25 |
| 18 | 47 | M | 133 | 92 | 67 | 12 | 6.5 | 14 |
| 19 | 49 | F | 105 | 59 | 95 | 12 | 10.5 | 10 |
| 20 | 64 | F | 125 | 85 | 70 | 12 | 9.7 | 17 |
| 21 | 78 | M | 122 | 80 | 92 | 12 | 15.3 | 19 |

Abbreviations: R<sub>o</sub> – resistance to cerebrospinal fluid outflow;
ICP – intracranial pressure;
ABPs – systolic arterial pressure;
ABPd – diastolic arterial pressure;
HR – heart rate;
BR – respiratory rate;
F – female; M – male.

### 3.2 Analysis of the original electrical model

Initial simulations based on previously described model parameters revealed several inconsistencies. In particular, the autoregulatory range of cerebral blood flow was narrower than expected under physiological conditions.

After modification of autoregulatory parameters, the simulated autoregulatory range was restored to the physiological interval of approximately 50–150 mmHg.

Further analysis revealed that some previously proposed electrical analog implementations underestimated arterial pulsation amplitudes. As a consequence, intracranial pressure pulsations in those models were unrealistically small. Correcting arterial pressure oscillations resulted in unrealistic intracranial pressure values during infusion test simulations, indicating the need for further modifications in the electrical analog.

### 3.3 Revised electrical analog model

To address these discrepancies, the electrical analog model was extended with additional components representing physiological damping of arterial pulsations transmitted to the cerebrospinal fluid compartment.

The full implementation of the revised model, including all additional components, is provided in Supplementary Fig. S1. The parameters of the electrical analog model, adopted from previous studies and modified in the present work, are summarized in Table 2.

**Table 2.**
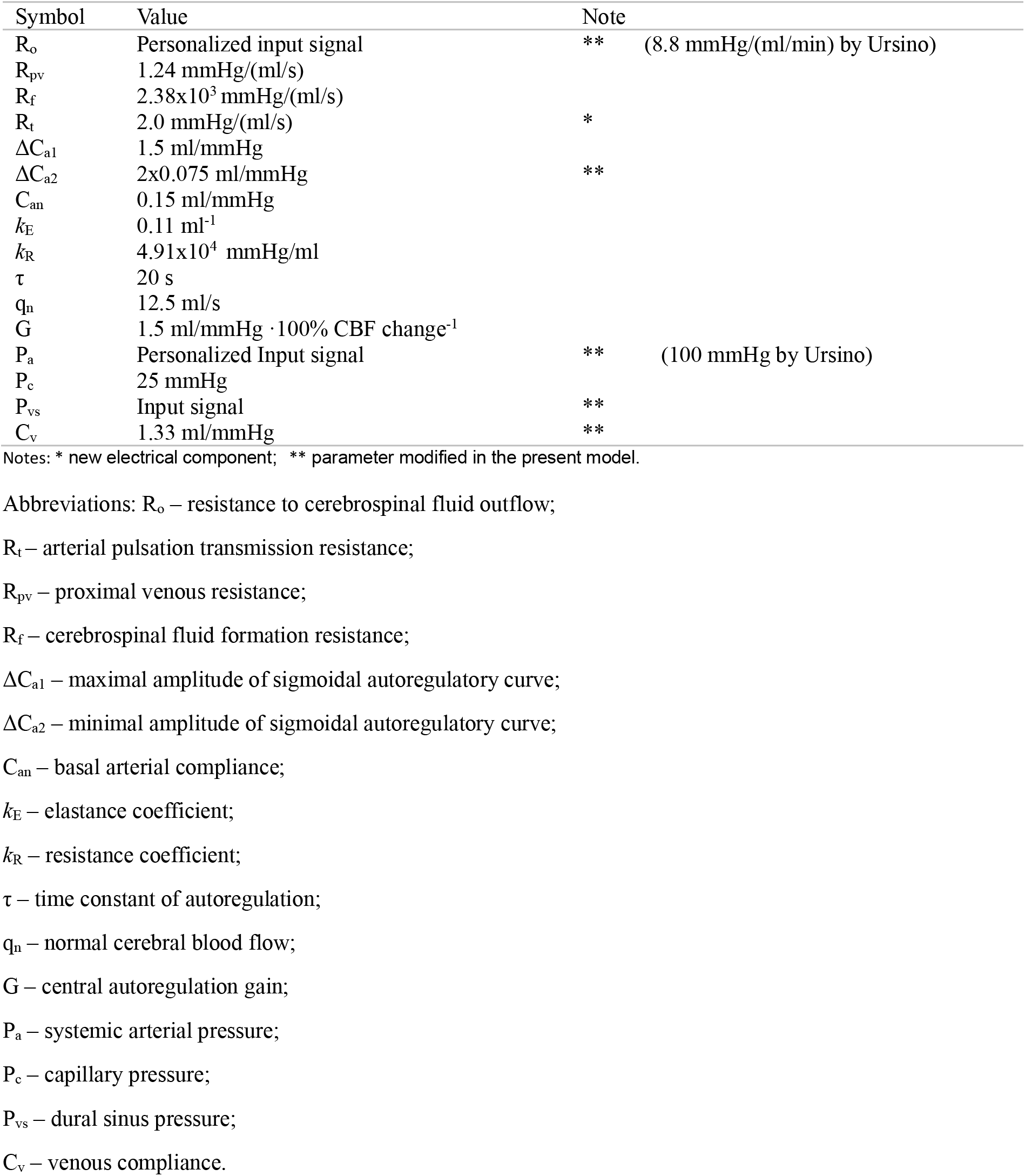
Parameters of the electrical analog model based on Ursino’s framework with modifications introduced in the present study.

### 3.4 Simulation of infusion tests

The modified model successfully reproduced the dynamic intracranial pressure changes observed during infusion tests. Fig. 3 presents an example of simulated intracranial pressure dynamics for patient case No. 19.

**Fig. 3.**
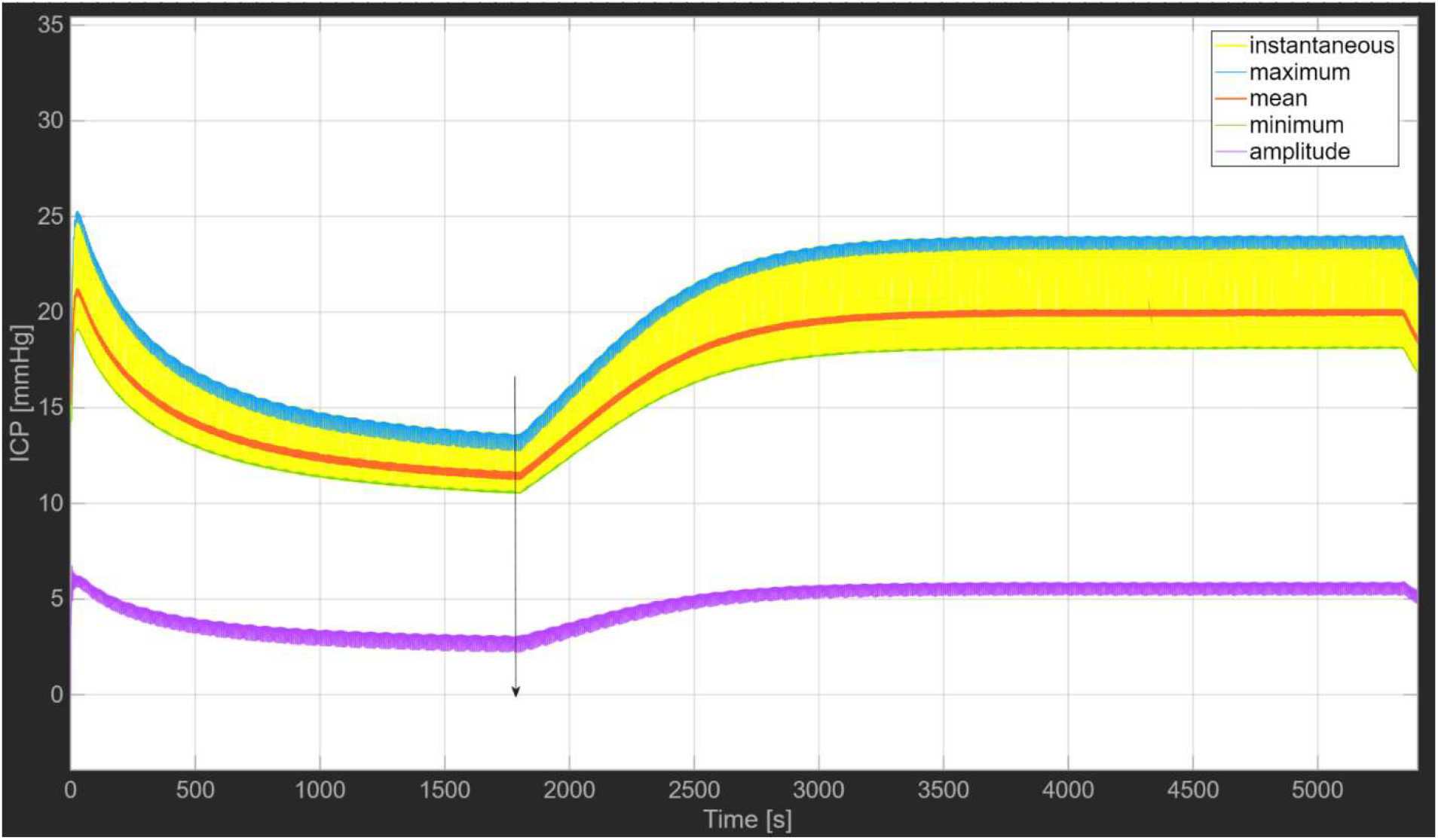
Simulated intracranial pressure dynamics during an infusion test. The figure illustrates simulated intracranial pressure (ICP) dynamics generated by the electrical analog model for patient case No. 19. After an initial stabilization period allowing the model to reach steady-state conditions, the simulated infusion results in a progressive rise of ICP followed by a plateau phase. After termination of the infusion, ICP gradually returns toward baseline values. The colored curves represent instantaneous ICP (yellow), maximum ICP (blue), mean ICP (red), minimum ICP (green), and pulse amplitude (purple). The progressive increase in pulse amplitude during infusion reflects the pressure-dependent reduction of intracranial compliance.

A stabilization period of 30 minutes was required before the simulated infusion test to allow the model to reach physiological equilibrium. The simulations also reproduced the physiological increase in ICP pulse amplitude accompanying the rise in mean intracranial pressure during the infusion phase.

### 3.5 Validation of simulated results

Comparison of simulated and measured intracranial pressure values demonstrated a strong agreement between the model and clinical recordings. For the representative case No. 19, the correlation coefficient between simulated and measured ICP values was r = 0.96 (Fig. 4).

**Fig. 4.**
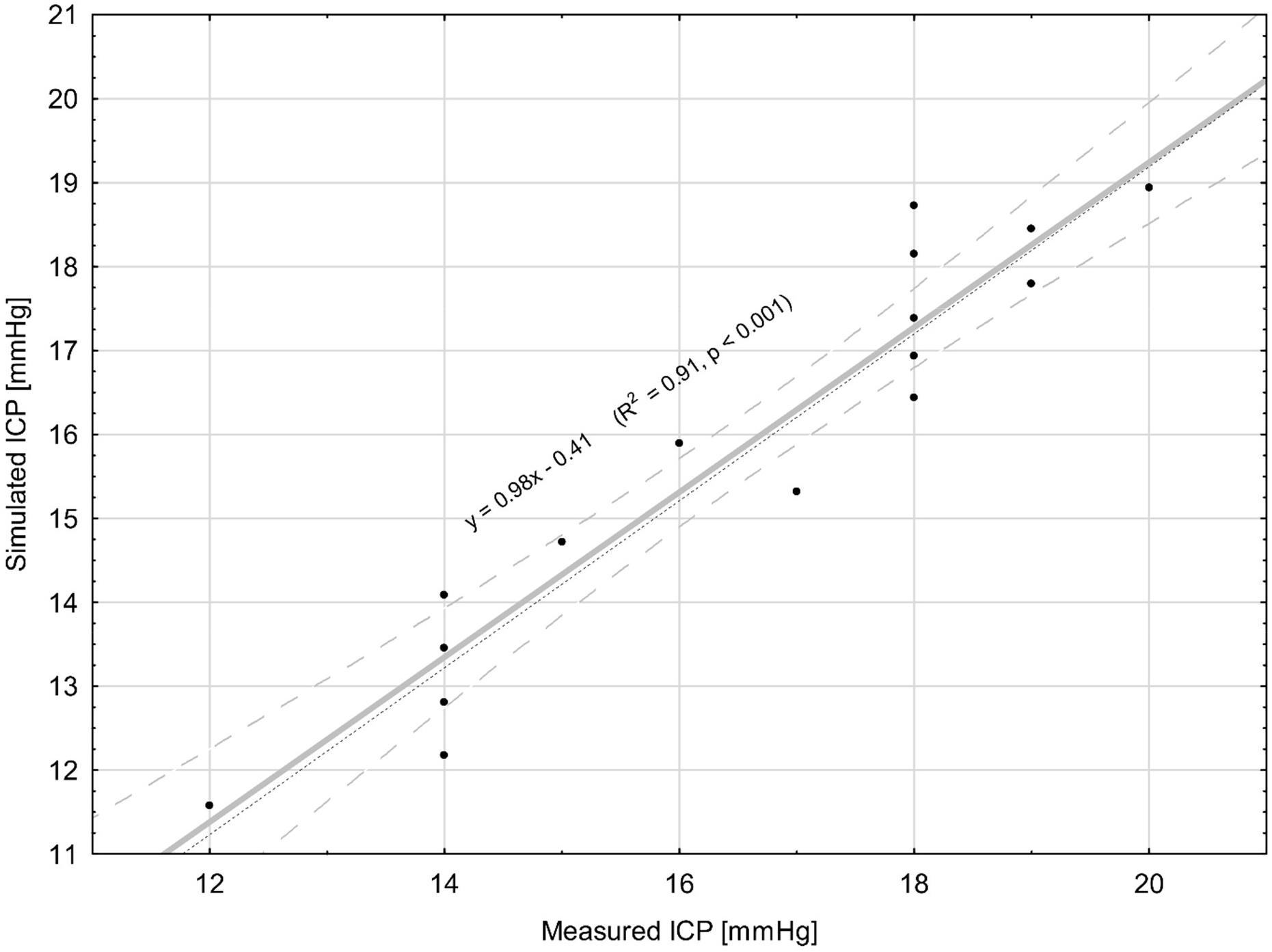
Linear regression between simulated and measured ICP values during infusion test (case 19). Linear regression analysis between simulated and measured intracranial pressure (ICP) values obtained during the cerebrospinal fluid infusion test in patient No. 19. Each point represents paired simulated and clinically recorded ICP values. The solid line indicates the linear regression fit (y = 0.98x - 0.41; R^2^ = 0.91, p < 0.001). The dashed line represents the line of identity (y = x). The shaded region denotes the 95% confidence interval of the regression model.

Graphical comparison of simulated and clinical ICP recordings demonstrated a close overlap between the curves (Fig. 5).

**Fig. 5.**
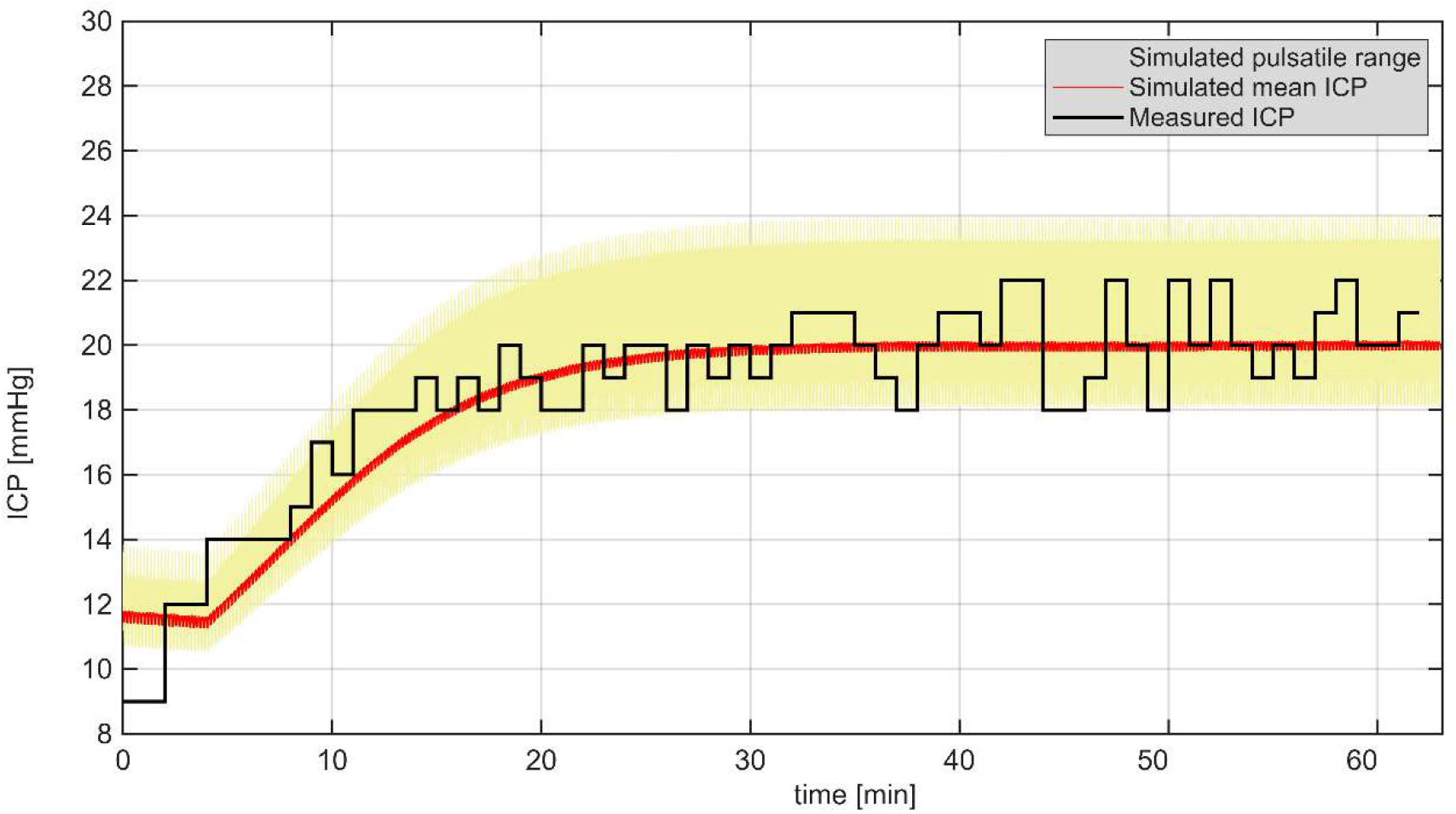
Comparison of simulated and clinical ICP recordings during infusion test. The graph shows the temporal evolution of intracranial pressure recorded during the clinical infusion test and the corresponding simulated ICP waveform generated by the electrical model. The yellow shaded region represents the pulsatile range (min–max) predicted by the model.

A total of 21 infusion test simulations were performed using patient-specific clinical data. Linear regression analysis demonstrated consistently high agreement between simulated and clinically measured intracranial pressure (ICP) values across all cases. The mean correlation coefficient (r) was 0.95 (95% CI: 0.94–0.96), with individual values ranging from 0.89 to 0.98 (Table 3).

**Table 3.** Comparison between simulated and measured ICP values across 21 infusion tests. The table summarizes correlation coefficients and residual ranges between simulated and clinically measured intracranial pressure values.

| Case | r | Residual range (mmHg) | RMSE (mmHg) |
| --- | --- | --- | --- |
| 1 | 0.95 | -1.6 to +0.8 mmHg | 2.62 |
| 2 | 0.95 | -3.0 to +3.0 mmHg | 2.97 |
| 3 | 0.93 | -0.9 to +0.8 mmHg | 2.02 |
| 4 | 0.97 | -1.3 to 1.05 mmHg | 3.84 |
| 5 | 0.92 | -1.20 to +1.10 mmHg | 2.06 |
| 6 | 0.97 | -0.80 to +1.4 mmHg | 1.48 |
| 7 | 0.93 | -1.80 to +1.25 mmHg | 1.81 |
| 8 | 0.95 | -0.85 to +0.80 mmHg | 1.72 |
| 9 | 0.96 | -1.40 to +1.20 mmHg | 2.01 |
| 10 | 0.89 | -2.20 to 1.90 mmHg | 1.35 |
| 11 | 0.94 | -1.05 to 1.20 mmHg | 2.51 |
| 12 | 0.96 | -0.75 to +0.90 mmHg | 1.30 |
| 13 | 0.94 | -1.40 to +2.25 mmHg | 1.68 |
| 14 | 0.92 | -5.0 to +4.0 mmHg | 2.74 |
| 15 | 0.97 | -2.60 to +1.50 mmHg | 1.63 |
| 16 | 0.94 | -0.90 to +0.55 mmHg | 2.24 |
| 17 | 0.98 | -1.25 to +2.50 mmHg | 2.63 |
| 18 | 0.92 | -2.80 to +2.90 mmHg | 2.14 |
| 19 | 0.96 | -1.30 to +0.95 mmHg | 0.98 |
| 20 | 0.95 | -2.10 to +2.70 mmHg | 1.48 |
| 21 | 0.96 | -1.80 to +2.45 mmHg | 2.26 |
| Mean | 0.95 | -1.71 to +1.68 mmHg | 2.07 |
| Minimum | 0.89 | -5.0 to +0.55 mmHg | 0.98 |
| Maximum | 0.98 | -0.75 to +4.0 mmHg | 3.84 |
| CI -95% | 0.94 | -2.17 to +1.25 mmHg | 1.77 |
| CI +95% | 0.96 | -1.26 to +2.10 mmHg | 2.37 |

Abbreviations: r – correlation coefficient; RMSE – root-mean-square error.

The corresponding mean coefficient of determination (R^2^) was 0.90, indicating that approximately 90% of the variance in measured ICP was explained by the model. Residual analysis was performed separately for each infusion test by determining the minimum and maximum differences between simulated and measured ICP values. Across the 21 cases, the mean of the minimum residuals was –1.71 mmHg, while the mean of the maximum residuals was +1.68 mmHg. These values indicate that, on average, the model deviation remained within approximately ±2 mmHg across the analyzed cohort.

The residual distributions were generally symmetrical and centered around zero, suggesting no systematic bias toward overestimation or underestimation.

The mean root-mean-square error between simulated and measured ICP values was 2.07 mmHg.

These results indicate that the electrical analog model reproduces clinical ICP dynamics during infusion tests with high accuracy.

## 4. Discussion

The simulations reproduced three fundamental features of intracranial pressure dynamics observed during clinical infusion tests: slow volume-driven pressure changes, cardiac-related pulsations, and the progressive increase in pulsatile amplitude associated with reduced intracranial compliance. The validation strategy combined three complementary levels of model assessment: waveform comparison, point-to-point regression analysis, and cohort-level statistical evaluation, providing a comprehensive evaluation of model performance.

The present study proposes an extended electrical analog model for the simulation of intracranial pressure (ICP) dynamics and demonstrates its validation using clinical infusion test data. The principal aim of this work was to develop a physiologically consistent framework capable of reproducing patient-specific ICP behavior and to verify its predictive ability under controlled clinical conditions. Unlike many previously proposed models, the present framework was directly validated against patient-specific clinical data obtained during standardized infusion tests.

Electrical analog modeling has long been used to represent physiological systems because of the mathematical equivalence between hydraulic and electrical circuits. In the context of intracranial dynamics, electrical analogs allow complex interactions between cerebral blood flow, cerebrospinal fluid (CSF) circulation, and intracranial compliance to be expressed in a compact mathematical form. However, many previously proposed models were primarily theoretical and were not directly validated against clinical measurements obtained under standardized conditions. The present work attempts to bridge this gap by integrating clinical infusion test data with a modified electrical analog implemented in a numerical simulation environment.

A key aspect of the proposed framework is the incorporation of patient-specific physiological inputs, including arterial blood pressure, heart rate, respiratory rhythm, and the resistance to CSF outflow determined from clinical infusion tests. By introducing these parameters into the electrical analog, the model can reproduce realistic ICP dynamics rather than generic theoretical waveforms. This approach allows the simulation to reflect the physiological variability observed in individual patients and provides a more clinically relevant representation of intracranial hydrodynamics.

The model was implemented in MATLAB/Simulink, which enabled dynamic simulations of intracranial processes and facilitated the analysis of transient responses during infusion tests. The simulation results demonstrated a strong agreement with clinical recordings obtained from 21 infusion tests. The mean correlation coefficient between simulated and measured ICP values reached r = 0.95, with relatively small residual differences between predicted and observed pressure values. Such a high level of agreement indicates that the model reproduces both the general trend and the temporal evolution of intracranial pressure changes during the infusion procedure.

An important observation from the simulations is the ability of the model to reproduce the characteristic phases of the infusion test, including the baseline stabilization period, the pressure rise during fluid infusion, the plateau phase, and the return toward baseline after termination of the infusion. The accurate reproduction of these phases suggests that the electrical analog captures the essential mechanisms governing intracranial pressure regulation. In particular, the interaction between arterial pulsatility, intracranial compliance, and CSF outflow resistance appears to be adequately represented within the proposed framework. The simulations reproduced the well-known physiological relationship between intracranial pressure and pulse amplitude, with increasing ICP associated with a progressive rise in pulsatile amplitude. This behavior reflects the nonlinear decrease in intracranial compliance as pressure increases and is consistent with observations from clinical infusion tests. The simulations also demonstrate that the increase in pulsatile ICP amplitude precedes the full rise of mean intracranial pressure during the infusion phase, reflecting the progressive reduction of intracranial compliance as intracranial volume increases. After termination of the infusion, mean intracranial pressure gradually decreases toward baseline values, while pulsatile amplitude remains transiently elevated. This behavior reflects the delayed recovery of intracranial compliance following transient intracranial volume loading and is consistent with clinical observations during infusion tests. Introduction of the additional resistance R_t_ proved necessary to limit excessive transmission of arterial pulsations to the cerebrospinal fluid compartment. In preliminary simulations performed without this element, restoration of physiologically realistic arterial pressure oscillations resulted in non-physiological ICP responses, including excessive pulse amplitudes and abnormally high plateau pressures. The R_t_ parameter therefore acts as a damping element enabling physiologically realistic coupling between arterial pulsatility and CSF pressure dynamics.

From a broader perspective, the proposed model can be interpreted as a computational representation - an “*electrical twin” -* of intracranial pressure dynamics under defined physiological conditions. Such a framework provides a useful platform for exploring how different physiological parameters influence ICP behavior and for testing hypotheses regarding the mechanisms underlying intracranial pressure regulation. Importantly, because the model can incorporate patient-specific parameters, it may serve as a tool for studying individual intracranial dynamics rather than only generalized theoretical scenarios. This multi-scale representation of intracranial pressure dynamics supports the interpretation of the model as a simplified electrical twin of intracranial hydrodynamics.

The results obtained in this study also highlight the potential clinical relevance of simulation-based approaches. Infusion tests remain an important diagnostic tool in the assessment of disorders related to CSF circulation, including hydrocephalus. However, interpretation of infusion test results may sometimes be challenging, particularly when complex interactions between vascular and CSF compartments are involved. Computational modeling may therefore provide an additional layer of analysis that helps interpret the physiological meaning of observed pressure changes.

Despite the promising results, several limitations of the present study should be acknowledged. First, the electrical analog remains a simplified representation of the intracranial system and cannot capture all aspects of cerebral physiology. Certain parameters were adopted from previously published models, while others were estimated or assumed due to the lack of direct experimental measurements. In particular, the additional resistance introduced between arterial and CSF compartments was assigned an approximate value because experimental data describing this parameter are not currently available. Second, the validation was performed using a limited number of infusion tests obtained under specific clinical conditions, and further studies involving larger datasets would be beneficial to confirm the general applicability of the model. Finally, the current implementation focuses on infusion test dynamics and does not yet incorporate other clinical scenarios, such as pathological alterations in cerebral autoregulation or long-term intracranial pressure variations.

Future work may therefore focus on refining the physiological representation of the model, incorporating additional experimental data, and extending the simulation framework to other clinical conditions. Such developments may further enhance the usefulness of computational modeling in the study of intracranial hydrodynamics.

## 5. Conclusions

This study introduces an extended electrical analog model for the simulation of intracranial pressure dynamics and demonstrates its validation using clinical infusion test data. By integrating patient-specific physiological inputs with a modified electrical circuit representation implemented in MATLAB/Simulink, the model is capable of reproducing realistic ICP behavior observed during clinical infusion tests.

The comparison between simulated and measured ICP recordings revealed a strong agreement, with a mean correlation coefficient of r = 0.95 across the analyzed cases. These results indicate that the proposed framework can reliably reproduce the temporal evolution of intracranial pressure during infusion procedures.

The developed model may therefore serve as a useful computational platform for studying intracranial hydrodynamics and exploring the interactions between vascular and cerebrospinal fluid compartments. In particular, it may support the interpretation of clinical infusion test results and contribute to the analysis of patient-specific intracranial pressure dynamics.

Further research involving larger clinical datasets and additional physiological parameters may help refine the model and expand its potential applications in both experimental and clinical settings.

## Author contributions

L.H. conceived the study, developed the model, performed simulations, analyzed the data, and wrote the manuscript.

## Data availability

The data supporting the findings of this study are available from the corresponding author upon reasonable request.

