## Supplementary figures and images for "A Patient-Specific Electrical Twin of Intracranial Pressure Dynamics Validated by Clinical Infusion Tests"

### Supplementary Figure S1

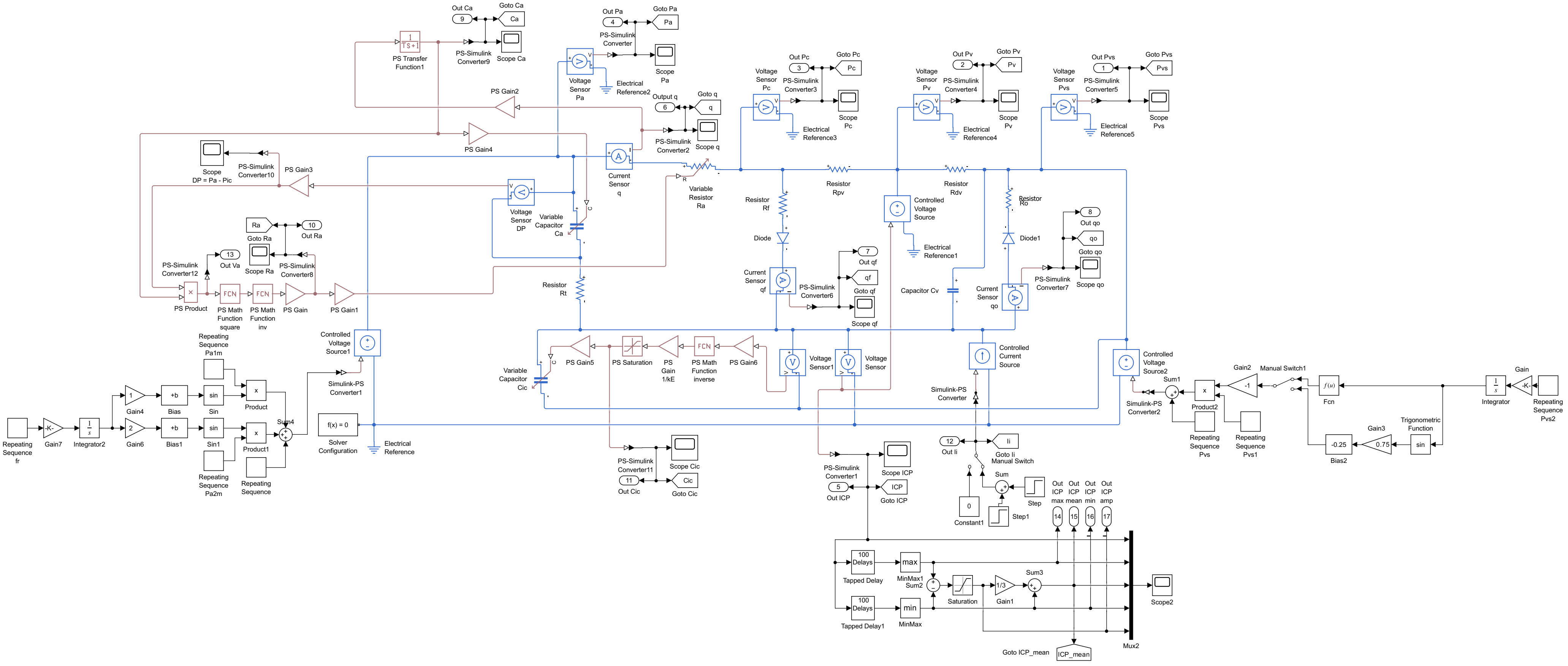
